# Leveraging host metabolism for bisdemethoxycurcumin production in *Pseudomonas putida*

**DOI:** 10.1101/753889

**Authors:** Matthew R. Incha, Mitchell G. Thompson, Jacquelyn M. Blake-Hedges, Yuzhong Liu, Allison N. Pearson, Matthias Schmidt, Jennifer W. Gin, Christopher J. Petzold, Adam M. Deutschbauer, Jay D. Keasling

## Abstract

*Pseudomonas putida* is a saprophytic bacterium with robust metabolisms and strong solvent tolerance making it an attractive host for metabolic engineering and bioremediation. Due to its diverse carbon metabolisms, its genome encodes an array of proteins and enzymes that can be readily applied to produce valuable products. In this work we sought to identify design principles and bottlenecks in the production of type III polyketide synthase (T3PKS)-derived compounds in *P. putida*. T3PKS products are widely used as nutraceuticals and medicines and often require aromatic starter units, such as coumaroyl-CoA, which is also an intermediate in the native coumarate catabolic pathway of *P. putida*. Using a randomly barcoded transposon mutant (RB-TnSeq) library, we assayed gene functions for a large portion of aromatic catabolism, confirmed known pathways, and proposed new annotations for two aromatic transporters. The 1,3,6,8-tetrahydroxynapthalene synthase of *Streptomyces coelicolor* (RppA), a microbial T3PKS, was then used to rapidly assay growth conditions for increased T3PKS product accumulation. The feruloyl/coumaroyl CoA synthetase (Fcs) of *P. putida* was used to supply coumaroyl-CoA for the curcuminoid synthase (CUS) of *Oryza sativa*, a plant T3PKS. We identified that accumulation of coumaroyl-CoA in this pathway results in extended growth lag times in *P. putida*. Deletion of the second step in coumarate catabolism, the enoyl-CoA hydratase-lyase (Ech), resulted in increased production of the type III polyketide bisdemethoxycurcumin.

## 1 INTRODUCTION

Secondary metabolites of fungi, plants, and bacteria have long been used as medicines and supplements (Hewlings and Kalman, 2017). Compounds such as naringenin, raspberry ketone, resveratrol, and curcumin are widely used nutraceuticals and are biosynthesized through similar pathways (Heller and Hahlbrock, 1980; Katsuyama et al., 2008; Schanz et al., 1992; Smith, 1996). Commercially, these chemicals are either extracted directly from plants or produced synthetically, as in the case of raspberry ketone (Smith, 1996). Renewable microbial production of these compounds will decrease reliance on agriculture and fossil fuel-derived chemical synthesis. The biosynthesis of these compounds (naringenin, raspberry ketone, resveratrol, and curcumin) relies on a class of enzymes called type III polyketide synthases (T3PKSs). T3PKSs carry out iterative Claisen condensation reactions typically with coenzyme A (CoA)-based starter and extender units (Yu et al., 2012). In the case of the 1,3,6,8-tetrahydroxynapthalene synthase of *Streptomyces coelicolor* (RppA) the starter and extender units are simply malonyl-CoA, while in many plant T3PKSs the starter unit is a phenylpropanoyl-CoA thioester, usually derived from ferulate, coumarate, or cinnamate (Izumikawa et al., 2003; Wakimoto et al., 2012).

Coumarate and ferulate are components of lignin found in lignocellulosic hydrolysate (LH), which has been proposed for use as a renewable feedstock for biocatalysis (Jönsson et al., 2013; Lawther and Sun, 1996). Characteristics such as high solvent tolerance and diverse carbon metabolisms are essential for microbes to be used in LH valorization. However, these robust traits are lacking in commonly used model organisms, such as *Escherichia coli*, hindering progress toward making LH a viable feedstock (Mills et al., 2009). Rather than developing these characteristics in a microorganism *de novo*, a clear alternative is to source a microbe with the desirable traits already available to it. The saprophytic bacterium *Pseudomonas putida* has long been studied for its ability to catabolize aromatic compounds and withstand solvents, making it an attractive host for LH upcycling (Guarnieri et al., 2017; Linger et al., 2014; Nozaki et al., 1963).

In this work we sought to leverage the native catabolism of *P. putida* KT2440 for use in the biosynthesis of a plant T3PKS product from coumarate. RppA of *S. coelicolor* was first expressed to determine production conditions conducive to T3PKS product accumulation. Using a randomly barcoded transposon mutant (RB-TnSeq) library of *P. putida*, we assayed for genes involved in the catabolism of coumarate and seven other related aromatic compounds. We then found that accumulation of coumaroyl-CoA by the activity of Fcs was toxic to *P. putida*. The native feruloyl/coumaroyl-CoA synthetase (Fcs) of *P. putida* was then used to produce the coumaroyl-CoA starter unit to the curcuminoid synthase (CUS) of *Oryza sativa*. Finally, relying on the native expression levels of *fcs* from the chromosome and plasmid-based expression of *CUS* (Figure 1), production of bisdemethoxycurcumin (BDC) from coumarate was achieved.

**Figure 1:**
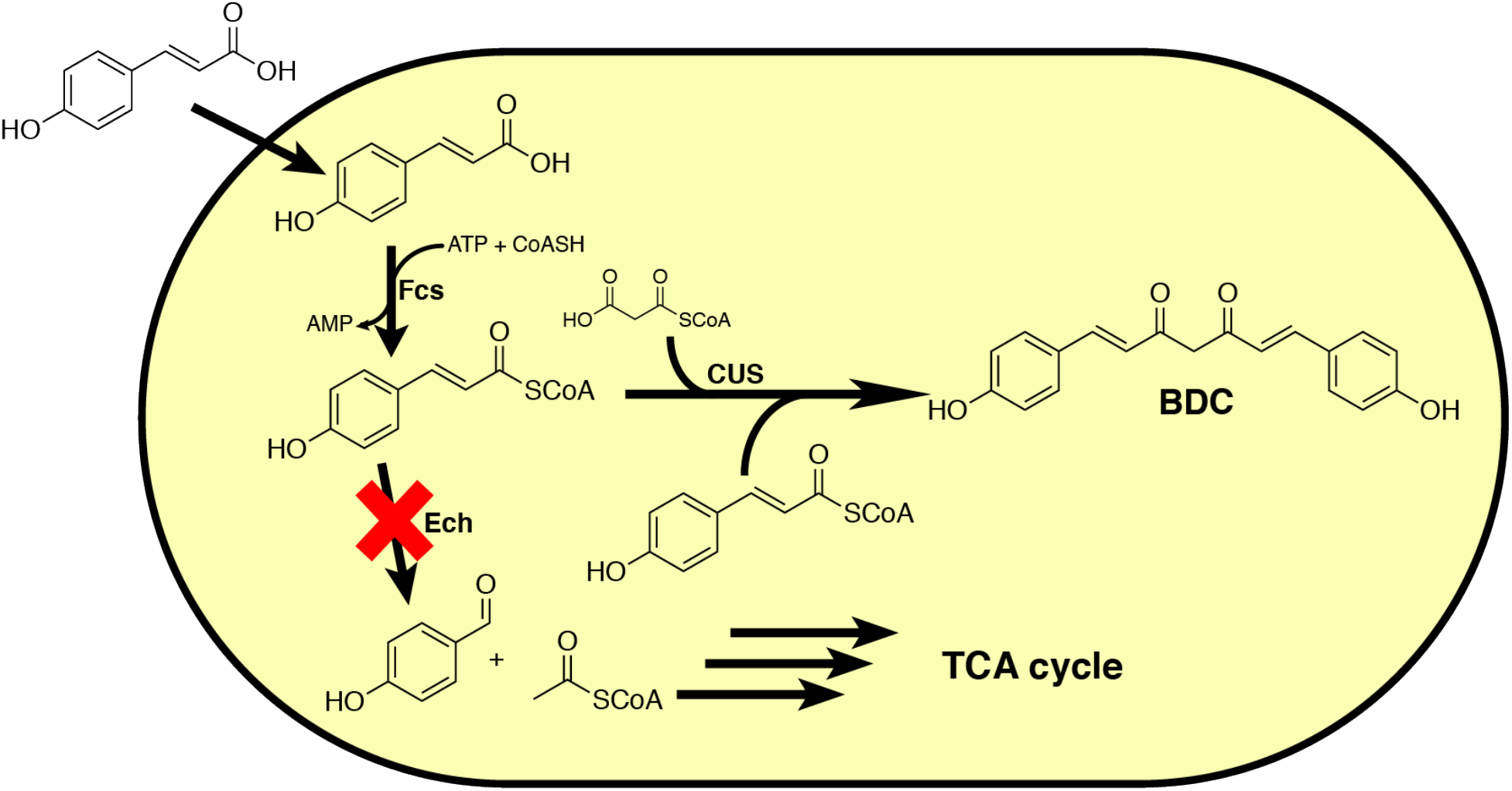
Diagram of the engineered *P. putida* strain used to produce bisdemethoxycurcumin. Fcs is the feruloyl/coumaroyl-CoA synthetase native to *P. putida* KT2440. Ech is the native enoyl-CoA hydratase-lyase which carries out the second step in coumarate catabolism and was knocked out in our production host. CUS is the curcuminoid synthase from *O. sativa.* BDC is the final curcuminoid product, bisdemethoxycurcumin.

## 2 RESULTS

### 2.1 RppA as a screen for optimal T3PKS production conditions

The tetrahydroxynaphthalene synthase of *Streptoyces coelicolor* was recently applied as a biosensor for malonyl-CoA concentrations in several hosts including *P. putida* (Yang et al., 2018). We sought to express this protein from the broad host range pBADT vector to assay culture conditions for increased product accumulation in *P. putida*. RppA was initially codon optimized for expression in *P. putida* and cloned into the construct pBADT-*rppA*-OW. We then constructed two more plasmids, one with the complete *rppA* cloned from the *S. coelicolor* genome (pBADT-*rppA*-NW) and another with a 3’ truncation of 75 base pairs (pBADT-*rppA*-NT) as this had been described previously to increase enzymatic activity (Izumikawa et al., 2003).

Since the product of RppA, 1,3,6,8-tetrahydroxynaphthalene, spontaneously oxidizes to the red pigment flaviolin, we used a colorimetric assay to determine how glucose concentrations affect the production of T3PKS products (Figure 2A). Concentrations of glucose ranging from 0-400 mM supplemented into LB medium were tested, and the production of flaviolin was measured by absorbance at 340 nm, as previously reported (Yang et al., 2018). Unexpectedly, the *rppA* expression vectors using the native *S. coelicolor* codons produced more flaviolin than the *P. putida* codon optimized variant (Figure 2B). In all three constructs tested, there was an increase in the accumulation of flaviolin in cultures containing greater than 25 mM glucose and accumulation of flaviolin appeared to plateau at ~100 mM glucose (Figures 2B and S4). To validate the observed red compound was flaviolin, we extracted, purified, and obtained an ^1^H-NMR spectrum for the red product using established methods (Figure S5A and S5B) (Gross et al., 2006; Soga, 1982). The final yield from 100 mL of culture was ~6.5 mg.

**Figure 2:**
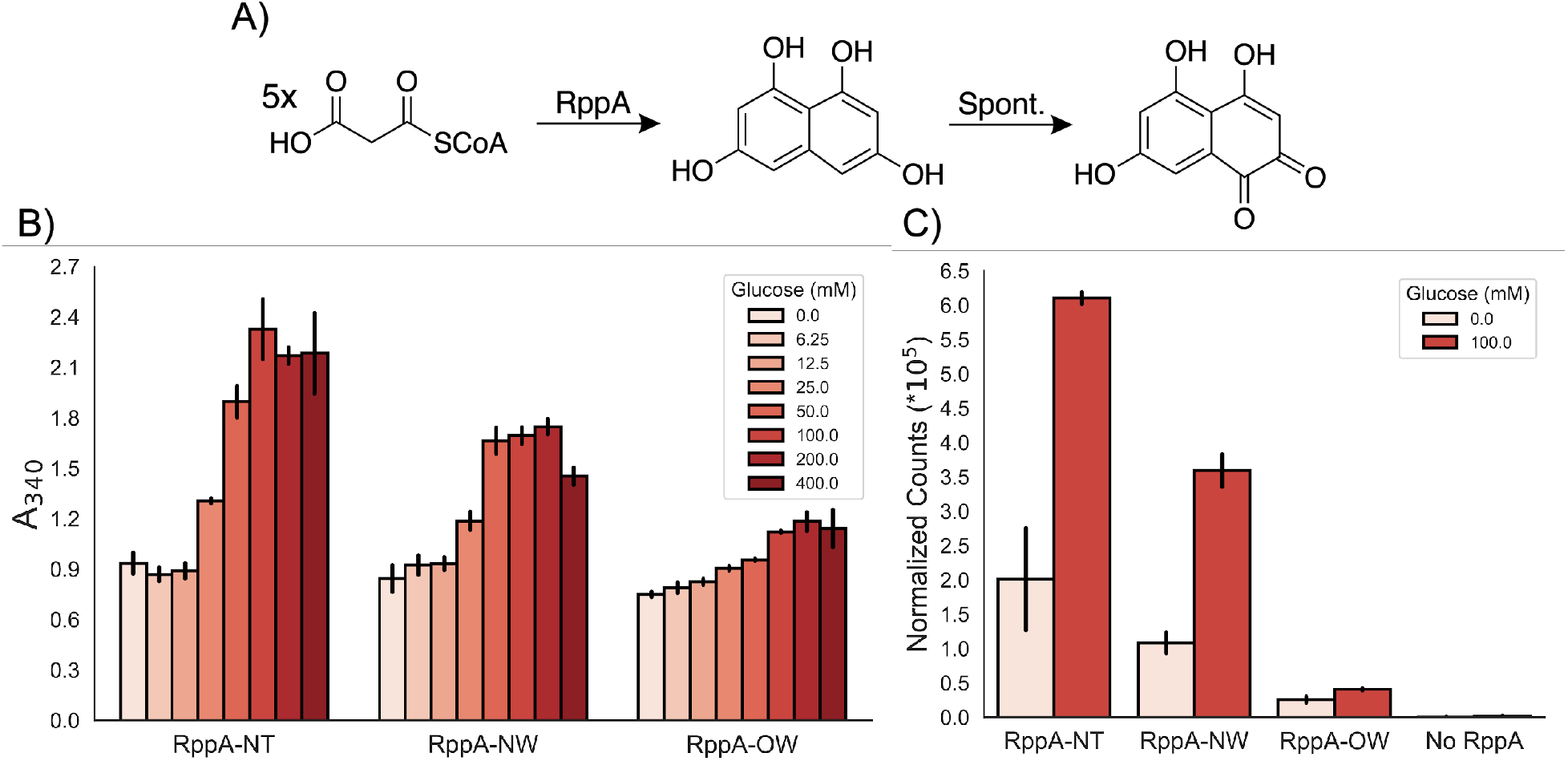
Flaviolin production as a reference for T3PKS activity. (A) Diagram of the pathway to the red pigment, flaviolin, from RppA and subsequent spontaneous oxidation. (B) Absorbance measurements at 340 nm of supernatants from cells expressing RppA variants in LB containing different concentrations of glucose. Error bars indicate the standard deviation of three trials. RppA-NW contains natural *S. coelicolor* codons, RppA-NT is the same as RppA-NW but with a C-terminal truncation of 25 amino acids, and RppA-OW is translated from a transcript that was codon optimized for *P. putida.* (C) RppA expression profile as determined by mass spectrometry from lysates of cultures producing the RppA variants. The No RppA sample was extracted from *P. putida* KT2440 carrying pBADT-RFP. Error bars indicate the standard deviation of three replicates.

Following this result, we sought to identify if increased protein abundance was the cause for increased flaviolin production. When RppA was quantified using LC-MS, we observed an increase in its relative abundance for all variants when 100 mM glucose was supplemented to the medium. The abundance of RppA also was highest in the strains expressing the native genomic sequence of *rppA* (Figure 2C).

### 2.2 Functional genomics to validate aromatic catabolisms of *P. putida*

While aromatic catabolism has been extensively studied in *P. putida*, essential genes implicated in these pathways have been described as recently as 2019 (Price et al., 2019). The genes involved in the first steps of coumarate catabolism reside in an operon with a putative acyl-CoA dehydrogenase (*PP_3354*) and a putative beta-ketothiolase (*PP_3355*), which have been proposed to be involved in an alternative catabolic pathway (Overhage et al., 1999). As functional redundancy in coumarate catabolism could result in loss of the type III polyketide precursor, coumaroyl-CoA, we sought to identify any pathways that could potentially impact product titers. To assay for genes involved in coumarate and related aromatic metabolisms, we grew a randomly barcoded transposon mutant (RB-TnSeq) library of *P. putida* KT2440 in minimal medium with a variety of different aromatic compounds often found in LH (p-coumarate, ferulate, benzoate, p-hydroxybenzoate, protocatechuate, vanillin, vanillate, phenylacetate) and glucose as sole carbon sources. The fitness of each gene was calculated by comparing the abundance of barcodes before versus after growth selection, using barcode sequencing (BarSeq) (Rand et al., 2017; Wetmore et al., 2015). Negative values indicate that the gene was important for growth in that condition.

The results of the RB-TnSeq assay validated that the primary route for ferulate and coumarate catabolism is through the feruloyl/coumaroyl-CoA synthetase (Fcs) and the enoyl-CoA hydratase lyase (Ech) (Figure 3 and Figure S1). The genes in the proposed secondary pathway of coumarate and ferulate catabolism, *PP_3354* and *PP_3355*, had no significant fitness phenotype, indicating that these genes are likely not necessary for coumarate or ferulate catabolism (Plaggenborg et al., 2003).

**Figure 3:**
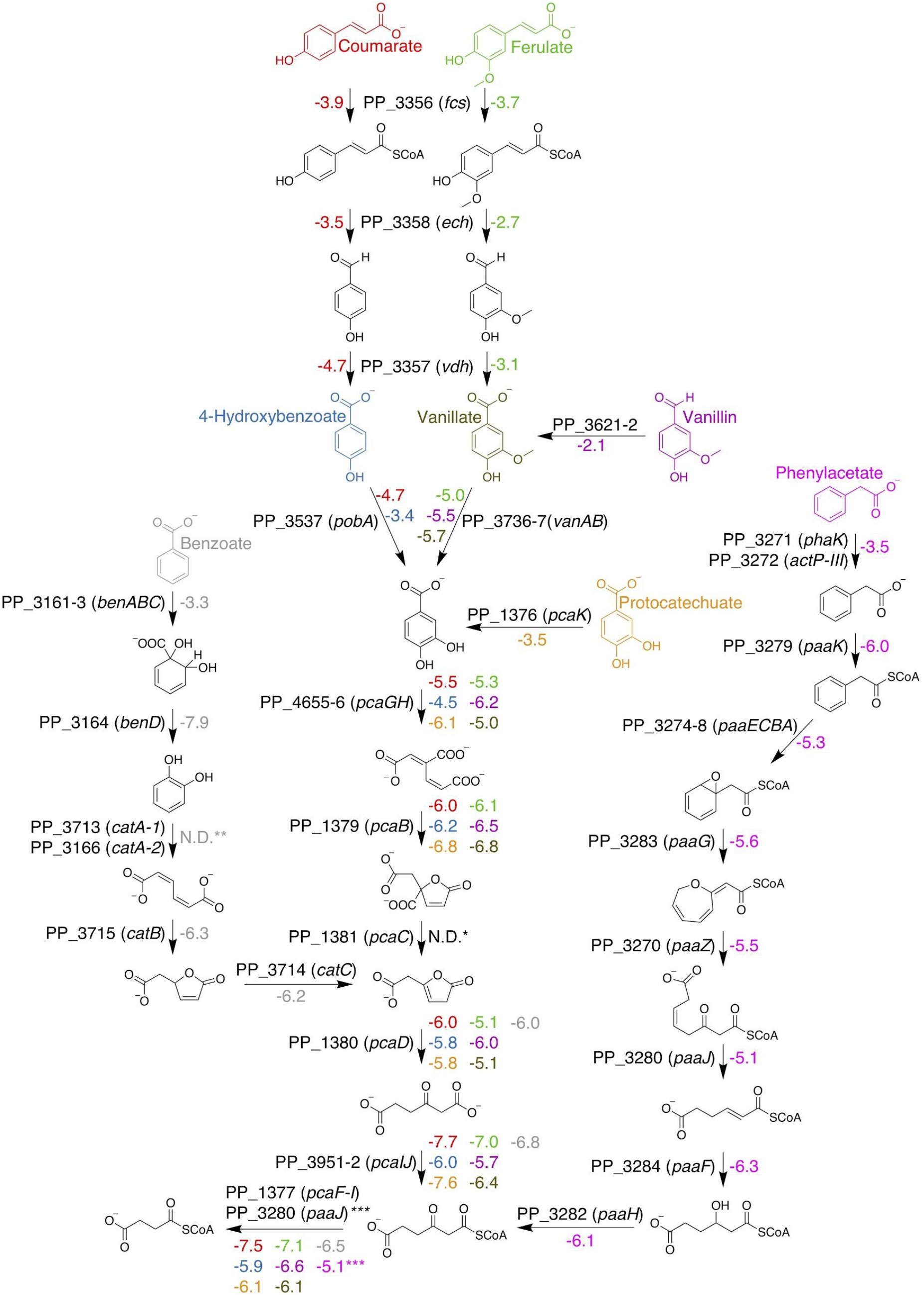
Overview of aromatic catabolism in *P. putida* KT2440. The colored and labelled compounds depicted were fed as sole carbon sources to the barcoded transposon library. Fitness values for each gene are the average of 2 replicate RB-TnSeq assays, and are colored corresponding to the carbon source (gray: benzoate, red: coumarate, blue: 4-hydroxybenzoate, light green: ferulate, dark green: vanillate, purple: vanillin, yellow: protocatechuate, pink: phenylacetate). For reactions where multiple genes are necessary, i.e. enzyme complexes, the fitness values for each gene involved in the reaction were averaged. *The t-score for *pcaC* was insignificant (|t_score_| < 4.0) and was excluded from our analysis. **Fitness values for these genes were mild (fitness > −2.0) and excluded. ***Fitness value corresponds to *paaJ*.

Of all the known reactions depicted in the map of aromatic catabolism (Figure 3), we observed significant negative fitness values for all but three of their corresponding genes: *catA-1*, *catA-2*, and *pcaC*. Even though the fitness score was higher than our cutoff at −1.4, *catA-1* had a significant |t-score| of 6.2 when grown on benzoate. This reflects previous work demonstrating CatA-I is the preferred catechol 1,2-dioxygenase, while CatA-II acts as a “safety valve” to handle high intracellular concentrations of catechol. It was also demonstrated that both catA-I and catA-II need to be deleted to abolish growth on benzoate, and this likely explains why the fitness value for catA-I was above our cut-off (Jiménez et al., 2014). In the case of *pcaC*, we noticed that there was a strong phenotype (fitness < −2.0), but the significance fell below our cutoff in all conditions tested (|t_score_| < 4.0). This is likely due to the low frequency of transposon insertions into this gene in the library (n=4).

While our results heavily support the current models of aromatic metabolism in *P. putida*, our data also indicated that some gene annotations should be revised. *PP_3272* is currently annotated as encoding an acetate permease (Kanehisa, 2019; Kanehisa et al., 2019; Kanehisa and Goto, 2000). We observed that *PP_3272* had a significant phenotype when grown on phenylacetate (Figure 3). Given these data and previously described homology to other systems (Jiménez et al., 2002), *PP_3272* should be reannotated as the phenylacetate transporter (phaJ). The *PP_1376* gene is annotated as encoding a 4-hydroxybenzoate transporter (Kanehisa, 2019; Kanehisa et al., 2019; Kanehisa and Goto, 2000); however, we only observed a fitness detriment for this gene with protocatechuate as the sole carbon source (Figure 3). Because of this, *PP_1376* should be reannotated as a protocatechuate transporter.

### 2.3 Accumulation of coumaroyl-CoA is toxic to *P. putida*

The first step in the biosynthetic pathway for bisdemethoxycurcumin is the activation of coumarate with coenzyme A (CoASH) (Figure 1). Because *Pseudomonas putida* KT2440 natively produces coumaroyl-CoA during coumarate catabolism, we knocked out the subsequent gene in the native catabolic pathway, *ech*, to prevent *P. putida* from consuming this necessary precursor (Figure 1) (Jiménez et al., 2002). Initial production experiments in Δ*ech* strains overexpressing *fcs* and *CUS* from a synthetic operon resulted in an extended lag phase (data not shown). To determine the cause, we overexpressed *fcs* alone under control of the arabinose-inducible *araBAD* promoter (P_BAD_) in the presence and absence of coumarate. Increasing the inducer concentration resulted in increased lag times only when coumarate was present in the medium (Figures 4A and S6). This suggested that the coumaroyl-CoA intermediate is toxic to *P. putida*.

**Figure 4:**
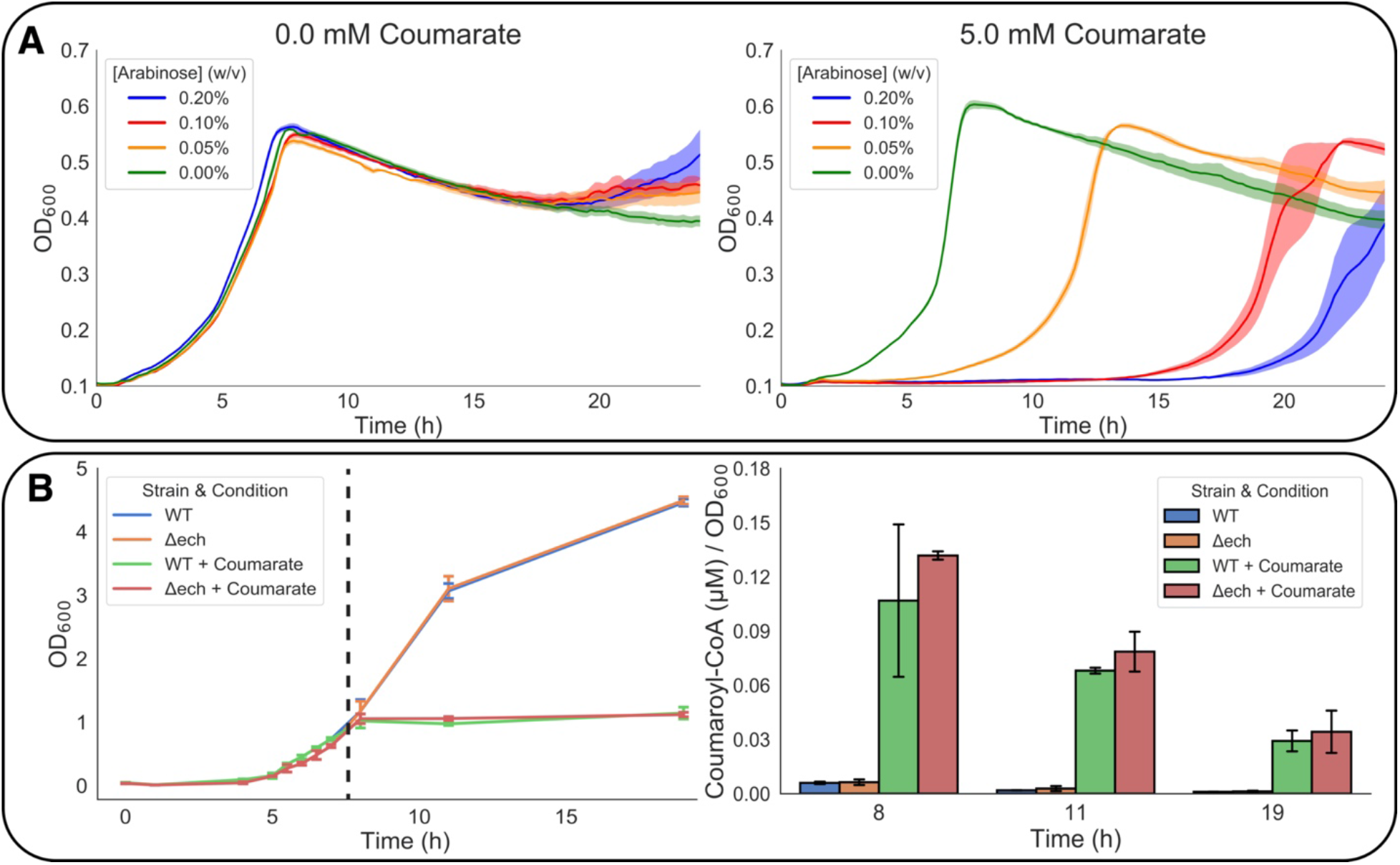
Production of coumaroyl-CoA inhibits growth of *P. putida.* (A) 24 hour growth curves of KT2440 Δ*ech* carrying pBADT-*fcs* in LB induced with varying concentrations of arabinose and no added coumarate (left), or 5 mM coumarate (right). Error bars are +/−the standard deviation of three replicates. (B) Growth curve (left) and normalized abundance of coumaroyl-CoA (right) in KT2440 (WT) and KT2440 Δ*ech* (Δ*ech*). Cultures expressing *fcs* were supplemented with 0 or 5 mM coumarate at hour 7 (marked with dashed line). Coumaroyl-CoA concentrations were normalized by OD_600_. Error bars are +/−the standard deviation of three replicates.

To determine if coumaroyl-CoA concentrations are elevated in cultures expressing *fcs*, cultures carrying pBADT-*fcs* were induced with L-arabinose. After a 7-hour growth period, 5 mM coumarate was supplemented to the medium. Following supplementation, growth was stunted and remained relatively constant over the course of 11 hours. The concentration of coumaroyl-CoA was highest in samples one hour after coumarate was supplemented into the medium; this was observed regardless of the *ech* genotype (Figure 4B). We therefore sought to minimize the burden of this intermediate by relying on the native chromosomal expression of *fcs* in future experiments.

### 2.4 Production of bisdemethoxycurcumin

In order to produce bisdemethoxycurcumin in *P. putida*, we constructed the biochemical pathway outlined in Figure 1. Exogenously added coumarate is activated by the native feruloyl/coumaroyl-CoA synthetase of *P. putida* (Fcs), then two resultant coumaroyl-CoA molecules are condensed with malonyl-CoA by the curcuminoid synthase of *O. sativa* (CUS) to yield bisdemethoxycurcumin (Katsuyama et al., 2007b). The CoA synthetase, *fcs*, was expressed from its native chromosomal locus, while *CUS* was expressed from the pBADT plasmid and induced with L-arabinose (Bi et al., 2013).

Our initial production strategy was to induce *CUS* until the cultures reached stationary phase. Then the cultures were pelleted and resuspended in fresh LB medium supplemented with 5 mM coumarate. These samples were then incubated for another 72 hours. A similar approach had been used successfully in *E. coli* (Katsuyama et al., 2007a); however, our titers were less than 0.5 mg/L (Figure S2). Given that bisdemethoxycurcumin is insoluble in water (Javeri and Chand, 2016), a 10 % v/v oleyl alcohol overlay was used to extract the product as the fermentation progressed. Production levels were low (approximately 0.1 mg/L) in *P. putida* Δ*ech* when the medium was supplemented with 10 mM coumarate, likely due to the toxicity of coumaroyl-CoA (Figure 5). Supplementation with 5 mM coumarate resulted in a ~5-fold increase in bisdemethoxycurcumin titers in the *P. putida* Δ*ech* strain relative to wild-type (Figure 5).

**Figure 5:**
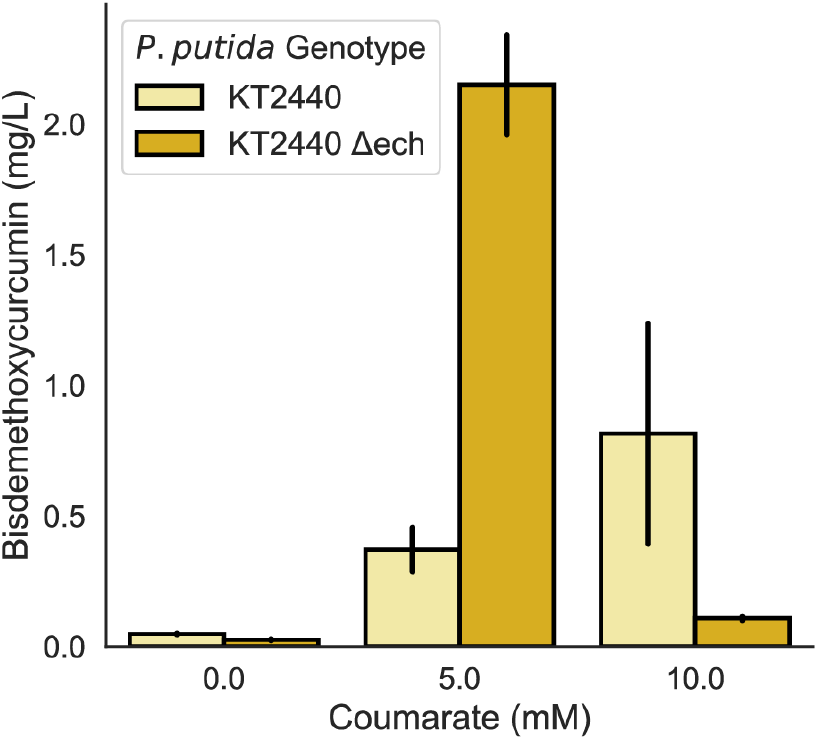
Bisdemethoxycurcumin titers in *P. putida* KT2440 Δ*ech* and in wildtype KT2440 harboring pBADT-CUS. *CUS* was induced with 2 % w/v L-arabinose. Error bars are +/−the standard deviation of three biological replicates.

## 3. DISCUSSION

*Pseudomonas putida* is among the most well studied saprophytic bacteria. Its diverse metabolisms enable it to catabolize a wide variety of complex carbon sources, including lignocellulosic hydrolysate (Wang et al., 2019). The robust catabolic pathways of *P. putida*, while useful for producing valuable molecules from diverse carbon sources, can also serve as an obstacle to achieving high product titers as it can often metabolize the desired products (Mitchell G Thompson et al., 2019). Given recent advances in gene editing techniques (Aparicio et al., 2019, 2018; Cook et al., 2018; Martínez-García and de Lorenzo, 2011; Wirth et al., 2019) and our ability to rapidly assay for gene function with transposon site sequencing (Price et al., 2018; Rand et al., 2017; Wetmore et al., 2015), engineering non-model hosts like *P. putida* for industrial applications has become less challenging.

Using RB-TnSeq mutant fitness assays we were able to rapidly confirm entire pathways of aromatic catabolism (Figure 3 and S1). The genes downstream of *fcs*, *PP_3354* and *PP_3355* previously described as a possible alternative route for coumarate/ferulate catabolism (Jiménez et al., 2002; Overhage et al., 1999; Priefert et al., 2001), showed no significant fitness detriment on any of the carbon sources tested. These genes may be structural remnants of a β-oxidation pathway that eventually evolved into the coumarate/ferulate pathway requiring *fcs*, *ech*, and *vdh* (Jiménez et al., 2002; Overhage et al., 1999; Priefert et al., 2001). We then proposed revised annotations of two genes required for transport of the aromatic compounds, phenylacetate (*PP_3272*) and protocatechuate (*PP_1376*), an important plant hormone and lignin metabolite respectively (Wightman and Lighty, 1982). There is a large amount of information in these data encompassing regulatory and structural genetic elements that could be useful to engineers and biologists.

Heterologous expression of bacterial T3PKSs, including the tetrahydroxynapthalene synthase of *S. coelicolor* (RppA), has previously been demonstrated in *P. putida* KT2440 (Gross et al., 2006; Yang et al., 2018). Using variants of *rppA*, we were able to rapidly screen type III polyketide production conditions. Expressing the codon optimized variant we created in this study, *rppA*-OW, resulted in less flaviolin production and less protein production than the native codon variants *rppA*-NW and *rppA*-NT. It is possible that there are some factors affecting heterologous protein expression that are not sufficiently accounted for in current codon optimization algorithms (Cambray et al., 2018). However, we demonstrated in all our constructs that increasing the glucose concentration had a considerable effect on the production of flaviolin (Figure 2 and S4).

Higher flaviolin accumulation with increased glucose concentrations could be due to increased flux to malonyl-CoA, a known limiting reactant to the biosynthesis of plant T3PKS products (Katsuyama et al., 2008; Zang et al., 2019), or due to increased functional protein expression. As the relative abundance of RppA increased when the culture was grown in the presence of 100 mM glucose, we hypothesize this likely contributed to the increased flaviolin titer observed in this condition (Figure 2B and 2C). The exact cause for the increase in protein abundance requires more detailed investigation. These T3PKS “sensors” have broad utility in both rapidly assaying culture conditions, as described here, and as high-throughput screens of genetic libraries for increased malonyl-CoA accumulation (Yang et al., 2018). Future work could employ these sensors to screen for increased intracellular malonyl-CoA concentrations from a complex growth medium like LH.

To provide the coumaroyl-CoA substrate for the bisdemethoxycurucumin T3PKS, CUS, we sought to use the native CoA synthetase (Fcs) of *P. putida*. Plasmid-based induction of *fcs* expression in the presence of coumarate, however, resulted in an increase in lag times due to the build-up of the toxic coumaroyl-CoA intermediate (Figures 4A, 4B, and S6). Neither the accumulation of coumaroyl-CoA nor the resultant growth inhibition seemed to be significantly affected by the genetic presence of *ech* (Figure 4B). The reduction in intracellular coumaroyl-CoA concentrations following the addition of coumarate was also not dependent on the presence of *ech* (Figure 4B). This reduction in thioester concentrations may be the result of nonspecific thioesterase activity, or spontaneous hydrolysis of coumaroyl-CoA.

The toxicity of hydroxycinnamate thioesters has been reported in *Acinetobacter baylyi* following disruptions in the gene encoding its enoyl-CoA hydratase lyase, and it was observed in *E. coli* expressing the *A. baylyi fcs* homolog in the presence of coumarate (Parke and Ornston, 2004). This toxicity could have been limiting other systems using bacterial coumaroyl-CoA synthetases, but the defective growth phenotype may not have been observed due to differences in experimental design (Park et al., 2009; Santos et al., 2011). The exact cause for coumaroyl-CoA toxicity remains unclear and will be the subject of future investigations.

To engineer *P. putida* for bisdemethoxycurcumin production, we deleted the native enoyl-CoA hydratase lyase (*ech*) responsible for the conversion of coumaroyl-CoA to acetyl-CoA and p-hydroxybenzaldehyde. In order to relieve some of the observed coumaroyl-CoA toxicity, we relied on the native genomic copy of *fcs* instead of a plasmid-based system. We demonstrated that native expression of *fcs* generates sufficient coumaroyl-CoA for curcuminoid synthase (CUS). Extraction of the product during growth using an oleyl alcohol overlay also significantly enhanced titers (Figure S2 and 5). In the final *P. putida* production strain, we achieved production of bisdemethoxycurcumin at titers of 2.15 mg/L (Figure 5).

This work is a significant first step towards the production of plant T3PKS-derived compounds in *P. putida*, but several issues remain to be addressed. In particular, our final bisdemthoxycurcumin titer of 2.15 mg/L is far lower than the 91.3 mg/l titer first described in *E. coli* (Katsuyama et al., 2008). Several methods have been proposed to enhance the production of curcuminoids and similar T3PKS derived compounds in microbial hosts. In one such approach, the metabolic burden of product, protein, and plasmid DNA synthesis was spread across several microbial strains in a synthetic co-culture (Fang et al., 2018). Since phenylpropanoid overproducing strains of *P. putida* have been described (Nijkamp et al., 2007, 2005), a co-culture method could be explored for the biosynthesis of plant T3PKS products in this host. It is also quite clear that more extensive pathway balancing will be required to mitigate the intermediate toxicity we have observed. As we continue to understand more about the metabolism of *P. putida* and learn to engineer it more effectively, this host will become an increasingly attractive chassis for renewable chemical biosynthesis.

## 4. METHODS

### 4.1 Media, chemicals, and culture conditions

General *E. coli* cultures were grown in Luria-Bertani (LB) Miller medium (BD Biosciences, USA) at 37 °C, while *P. putida* was grown at 30 °C. MOPS minimal medium was used where indicated and comprised of the following: 32.5 μM CaCl_2_, 0.29 mM K_2_SO_4_, 1.32 mM K_2_HPO_4_, 8 μM FeCl_2_, 40 mM MOPS, 4 mM tricine, 0.01 mM FeSO_4_, 9.52 mM NH_4_Cl, 0.52 mM MgCl_2_, 50 mM NaCl, 0.03 μM (NH_4_)_6_Mo_7_O_24_, 4 μM H_3_BO_3_, 0.3 μM CoCl_2_, 0.1 μM CuSO_4_, 0.8 μM MnCl_2_, and 0.1 μM ZnSO_4_ (LaBauve and Wargo, 2012). Cultures were supplemented with kanamycin (50 mg/L, Sigma Aldrich, USA) when indicated. Technical grade oleyl alcohol was acquired from Alfa Aesar (Alfa Aesar, Thermo Fisher Scientific). Coenzyme A (lithium salt) was purchased from CoALA Biosciences (CoALA Biosciences, USA). All other compounds were purchased through Sigma Aldrich (Sigma Aldrich, USA). Construction of *P. putida* deletion mutants was performed as described previously (Thompson et al., 2019).

### 4.2 Strains and plasmids

Bacterial strains and plasmids used in this work are listed in Table 1. All strains and plasmids created in this work are available through the public instance of the JBEI registry https://public-registry.jbei.org. All plasmids were designed using Device Editor and Vector Editor software, while all primers used for the construction of plasmids were designed using j5 software (Chen et al., 2012; Ham et al., 2012; Hillson et al., 2012). Plasmids were assembled via Gibson Assembly using standard protocols (Gibson et al., 2009), or Golden Gate Assembly using standard protocols (Engler et al., 2008). Plasmids were routinely isolated using the Qiaprep Spin Miniprep kit (Qiagen, USA), and all primers were purchased from Integrated DNA Technologies (IDT, Coralville, IA).

**Table 1:** Strains and plasmids used in this study.

| Strain | JBEI Part ID | Reference |
| --- | --- | --- |
| *E. coli* DH10B |  | (Grant et al., 1990) |
| *E. coli* BL21(DE3) |  | Novagen |
| *P. putida* KT2440 |  | ATCC 47054 |
| *P. putida* KT2440 $\Delta ech$ | | |
| **Plasmids** |  |  |
| pET28 |  | Novagen |
| pBADT |  | (Bi et al., 2013) |
| pBADT-*CUS* | JBx_134311 | This work |
| pBADT-*rppA*-NT | JBx_134312 | This work |
| pBADT-*rppA*-NW | JBx_134313 | This work |
| pBADT-*rppA*-OW | JBx_134314 | This work |
| pBADT-*fcs* | JBx_134315 | This work |
| pBADT-*fcs-CUS* | JBx_134316 | This work |
| pMQ30 |  | (Shanks et al., 2009) |
| pMQ30-*PP_3358* |  | (Park et al., 2019) |

### 4.3 Plate based growth assays

Growth studies of bacterial strains were conducted using microplate reader kinetic assays. Overnight cultures were inoculated into 10 mL of LB medium from single colonies, and grown at 30 °C. These cultures were then diluted 1:100 into 500 μL of LB medium with appropriate concentrations of arabinose and p-coumarate in 48-well plates (Falcon, 353072). Plates were sealed with a gas-permeable microplate adhesive film (VWR, USA), and optical density and fluorescence were monitored for 48 hours in a Biotek Synergy 4 plate reader (BioTek, USA) at 30 °C with fast continuous shaking. Optical density was measured at 600 nm. The amount of time necessary for the culture to reach an OD_600_ of 0.16 was defined as the lag time.

### 4.4 HPLC detection of bisdemethoxycurcumin

HPLC analysis was performed on an Agilent Technologies 1200 series liquid chromatography instrument coupled to a Diode Array Detector (Agilent Technologies, USA). Compounds were separated at a constant flow rate of 0.4 mL/min over a Kinetex C18 column (2.6 μm diameter, 100Å particle size, dimensions 100 × 3.00 mm, Phenomenex, USA) held at 50°C. The mobile phase consisted of H_2_O + 0.1% trifluoroacetic acid (A) and acetonitrile + 0.1% trifluoroacetic acid (B). Separation was performed using the following gradient method: 0-3 minutes 95% A, 3-15 minutes 95-5% A, 15-17 minutes 5% A, 17-17.5 minutes 5-95% A, 17.5-20 minutes 95% A. The presence of bisdemethoxycurcumin was monitored and quantified at 440 nm.

### 4.5 RB-TnSeq experiments and analysis

BarSeq-based experiments utilized the *P. putida* RB-TnSeq library, JBEI-1, which has been described previously (Thompson et al., 2019). An aliquot of JBEI-1 was thawed on ice, diluted into 25 mL of LB medium supplemented with kanamycin and grown to an OD_600_ of 0.5 at 30 °C. Three 1 mL aliquots of the library were pelleted and stored at −80 °C to later serve as the t_0_ of gene abundance. Libraries were then washed in MOPS minimal medium and diluted 1:50 in MOPS minimal medium with 10 mM p-coumarate, ferulate, benzoate, p-hydroxybenzoate, protocatechuate, vanillin, vanillate, phenylacetate, or D-glucose. Cells were grown in 600 μL of medium in 96-well deep well plates (VWR). Plates were sealed with a gas-permeable microplate adhesive film (VWR, USA), and then grown at 30 °C in an INFORS HT Multitron (Infors USA Inc.), with shaking at 700 rpm. Two 600-μL samples were combined, pelleted, and stored at −80 °C until analysis by BarSeq, which was performed as previously described (Rand et al., 2017; Wetmore et al., 2015). All fitness data are publicly available at http://fit.genomics.lbl.gov.

### 4.5 Curcuminoid production

For production of bisdemthoxycurcumin without an overlay, an overnight culture of *P. putida* KT2440 Δ*ech* + pBADT-CUS was diluted 1:100 into 5 mL of LB supplemented with 50 mg/L kanamycin, 1 % w/v L-arabinose, and 100 mM glucose. The culture was grown to stationary phase over 12 hours then pelleted in a centrifuge at 5000 xg for 5 minutes. The cell pellets were resuspended in 2 mL of fresh LB with 50 mg/L kanamycin, 0.5 % L-arabinose, 100 mM glucose, and 5 mM coumarate. The culture was allowed to proceed for 72 hours.

For experiments employing an overlay, overnights of *P. putida* harboring pBADT-CUS were diluted 1:100 into 25 mL of fresh LB supplemented with 50 mg/L kanamycin, and 100 mM glucose. Arabinose and coumarate were added at the beginning of the fermentation at concentrations indicated. A 2.5 mL overlay of oleyl alcohol was added to extract the bisdemethoxycurcumin during growth. The fermentation was allowed to proceed for 72 hours.

### 4.6 Curcuminoid extraction

For cultures lacking an overlay, 0.5 mL of culture was acidified to pH 3 with 3 N HCl. Bisdemethoxycurcumin was then extracted with an equal volume of ethyl acetate. 250 μL of the ethyl acetate layer was removed and the solvent was allowed to evaporate overnight. The dried samples were then resuspended in 50 μL of acetonitrile for analysis with HPLC-DAD.

For cultures with an overlay, the cultures were acidified to pH 3 with 3 N HCl. Acidified cultures were then pelleted in a centrifuge and the oleyl alcohol overlays were removed. To quantify bisdemethoxycurcumin, 100 μL of the extracted overlays were added to a black, clear bottom 96-well plate and absorbance was measured at 425 nm in a Biotek Synergy 4 plate reader (BioTek, USA). A standard curve was made with bisdemethoxycurcumin standards dissolved in oleyl alcohol (Figure S3)

### 4.7 Flaviolin production and targeted proteomic analysis

Colonies of *P. putida* KT2440 strains carrying pBADT-*rppA*-NW, pBADT-*rppA*-NT, and pBADT-*rppA*-OW were used to inoculate LB or MOPS minimal medium with 50 mg/L kanamycin and cultured overnight. The overnight culture was then diluted 1/100 into fresh medium with 0.2 % w/v L-arabinose, 50 mg/L kanamycin, and 400, 200, 100, 50, 25, 12.5, 6.25 or 0 mM glucose. Cultures were conducted in 24-well deep-well plates and allowed to proceed for 48 hours. Cultures were then pelleted in a centrifuge at 5000 xg for 5 minutes. Supernatants were removed, aliquoted into a 96-well black clear bottom plate, and the absorbance was measured at 340 nm in a Biotek Synergy 4 plate reader (BioTek, USA). For cultures expressing *rppA* variants in LB, the supernatants were too opaque with red product and needed to be diluted 1:5 in fresh LB for accurate absorbance measurements. Supernatants of strains cultured in minimal medium were not diluted before absorbance measurements.

Samples for intracellular targeted proteomic analysis were grown for 48 hours in 5 mL LB supplemented with 0 mM or 100 mM glucose. Then, 2 mL of the culture were then pelleted by centrifugation, the supernatant was decanted, and the pellets were stored at −80 °C. Proteins were extracted and analyzed using a variation of a previously-described workflow (Chen et al., 2019).

### 4.8 Flaviolin purification

An overnight culture of *P. putida* KT2440 carrying pBADT-rppA-NT was diluted 1:100 into 100 mL of LB supplemented with 25 mg/L kanamycin, 0.2% *w*/*v L*-arabinose, and 100 mM glucose. The culture was shaken for 48 hours at 30 °C in a 500 mL baffled Erlenmeyer flask. Following fermentation, the red supernatant was removed, acidified with 10 mL 10 *N* HCl, and extracted twice with equal volumes of ethyl acetate. The combined organic phase was evaporated leaving a black solid, which was then dissolved in ethyl acetate and methanol (7:3). The extract components were then separated using preparative thin layer chromatography (TLC, silica gel). The red band was isolated from the TLC plate, extracted with methanol, and filtered. The methanol was evaporated *in vacuo* leaving dark red solid with a yield of ~6.5 mg. The identity of the flaviolin was then validated with ^1^H-NMR. ^1^H-NMR (400 MHz, CD_3_OD) *δ* 6.99 (s, 1 H), 6.47 (s, 1 H), 5.58 (s, 1 H) (Figure S5A and S5B).

### 4.9 Synthesis of coumaroyl-CoA standard

**Scheme 1.**
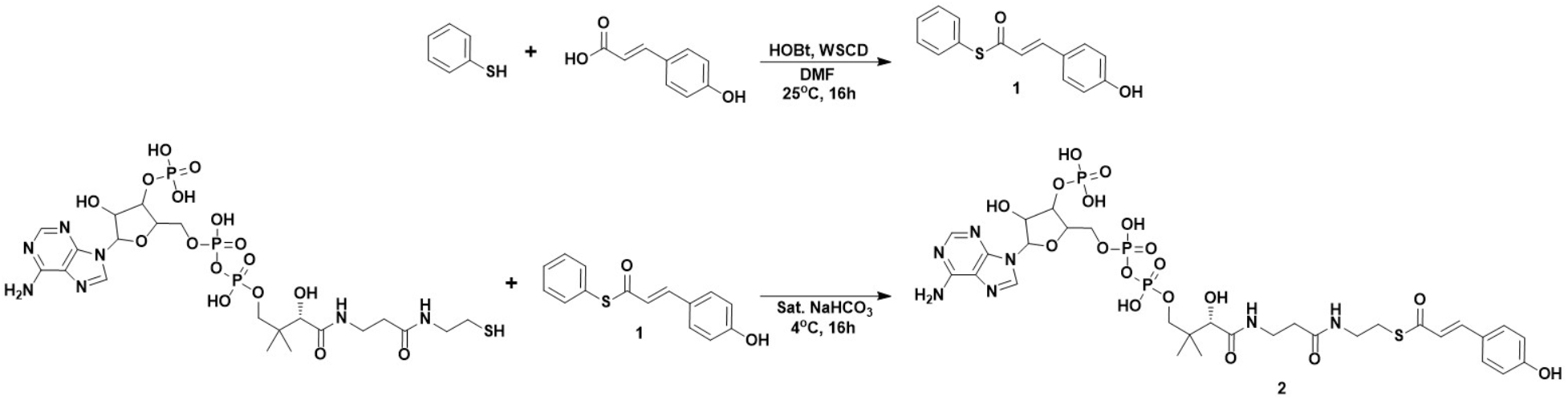

Coumaroyl-Coenzyme A (**2**) was synthesized in two steps according to Scheme 1. *S*-phenyl (*E*)-3-(4-hydroxyphenyl)prop-2-enethioate (**1**) was synthesized according to previous literature (Hori et al., 2012). 1-(3-Dimethylaminopropyl)-3-ethylcarbodiimide hydrochloride (WSCD·HCl, 1.48 g, 7.3 mmol), 1-hydroxybenzotriazole (HOBt, 1.12 g, 7.3 mmol), and 4-hydroxycinnamic acid (1 g, 6.1 mmol) were dissolved in DMF (100 mL) and stirred for 1 h at room temperature. Thiophenol (0.610 mL, 6.1 mmol) was subsequently added to the mixture and stirred overnight at room temperature. After the solvent was removed *in vacuo*, water and dichloromethane (DCM) were added to the mixture. The organic layers were combined, dried over Na_2_SO_4_, and evaporated. The product was purified with silica-gel column chromatography and eluted with DCM to yield *S*-phenyl (*E*)-3-(4-hydroxyphenyl)prop-2-enethioate as a white solid (**1**, 468 mg, 30%).

To synthesize coumaroyl-CoA (**2**), coenzyme A lithium salt (7.7 mg, 10 μmol) was dissolved in a solution of saturated sodium bicarbonate (750 μL) which had been pre-chilled to 4°C. *S*-phenyl (*E*)-3-(4-hydroxyphenyl)prop-2-enethioate (30 mg, 117 μmol) was added to the solution. The solution was mixed for 16 hours at 4 °C then quenched via the addition of 6M HCl to bring the pH of the solution to ~2. The mixture was extracted with ethyl acetate (2 × 2 mL), then coumaroyl-CoA was purified from the remaining aqueous phase via preparatory HPLC.

Coumaroyl-CoA was purified using an Agilent 1260 series preparatory HPLC system equipped with a 900 μL sample loop and a UV detector coupled to a fraction collector. Compounds were separated over an Agilent Zorbax SB-C18 preparatory column (9.4 × 50 mm, 5μm particle size) using a mobile phase composed of 10 mM ammonium formate, pH 4.5 (solvent A) and methanol (solvent B) using the following method:

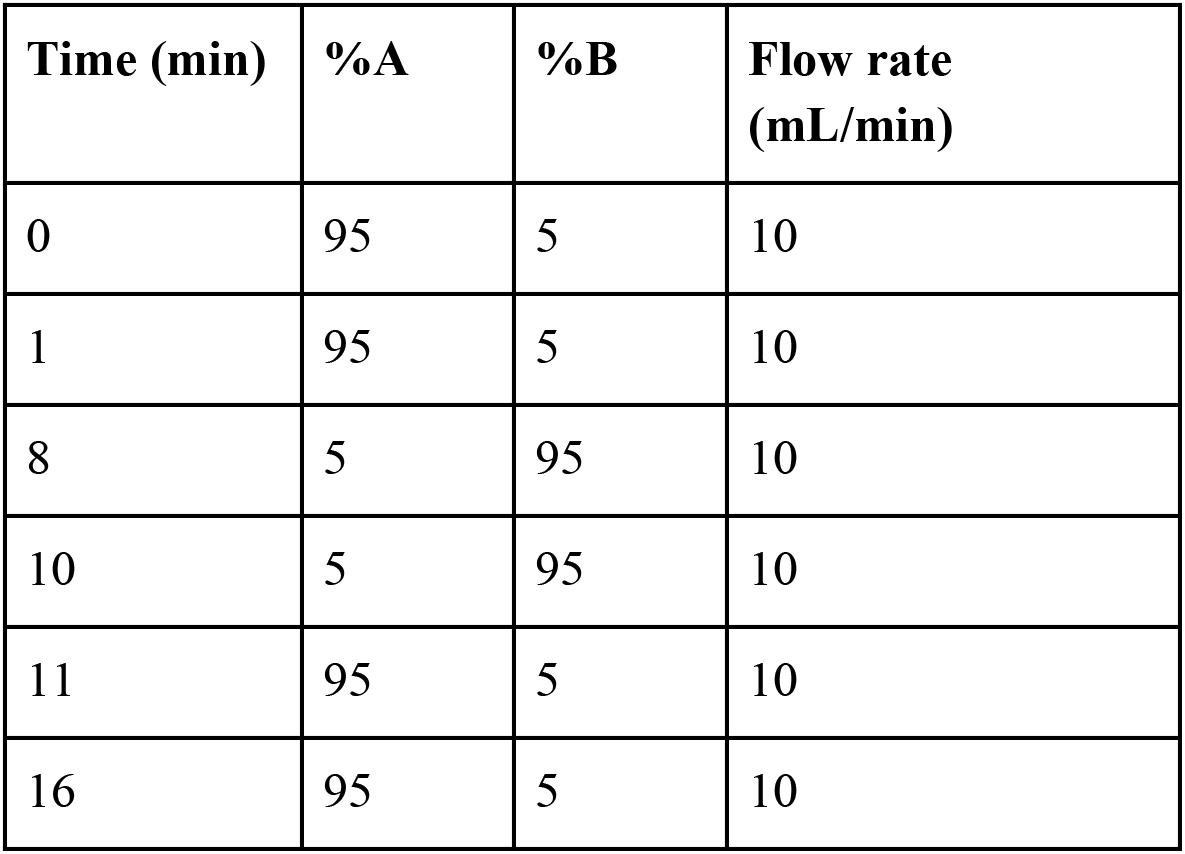

The fraction collector was programmed to collect peaks absorbing at a wavelength of 260 nm and that eluted between 3.6 and 4.6 minutes. The fractions containing coumaroyl-CoA were identified by direct infusion onto an Applied Biosystems 4000 QTRAP mass spectrometer system operating with the following parameters: Q1 MS mode, scan range 100-1000 m/z, declustering potential: 70, entrance potential: 10, curtain gas: 10, IonSpray Voltage: 4800, Temperature: 300, Ion Source Gases: 40. Fractions containing coumaroyl-CoA were combined and lyophilized, yielding coumaroyl-CoA as a white powder (6 mg,

### 4.10 LC-MS/MS analysis of intracellular Coenzyme A esters

*P. putida* KT2440 and *P. putida* Δ*ech* cells carrying pBADT-*fcs* were cultured overnight. Cultures were inoculated 1:100 into 25 mL LB supplemented with 100 mM glucose and 0.2% w/v L-arabinose. Cells were cultured at 30 °C for 7 hours then coumarate was added to a final concentration of 5 mM. 10 mL of culture were taken at hours 8, 11, and 19 and pelleted in a centrifuge. Cell pellets were resuspended in 1 mL ice cold methanol and stored at −80 °C until workup.

Coenzyme A esters were extracted as follows: to the 1 mL methanol resuspensions, 0.5 mL of ice cold LC grade water and 0.25 mL ice cold LC chloroform. Samples were vortexed to mix after each addition. Samples were then centrifuged at maximum speed for 2 minutes to promote phase separation, after which 0.5 mL of the top aqueous phase was removed and transferred to an Amicon 3 kDa molecular weight cutoff centrifuge filter (Sigma Aldrich, USA). Extracts were filtered by spinning in a benchtop centrifuge at 4 °C for 90 minutes at 13000g. The flow through was lyophilized for 24 hours to concentrate the sample. Lyophilized fractions were resuspended in 0.1 mL of ice cold 60:40 (v/v) LC grade acetonitrile: LC grade water, and 35 μL of the aqueous layer was removed for LC-MS/MS analysis.

CoA extracts were analyzed using an Applied Biosystems 4000 QTRAP mass spectrometer coupled to an Aglient 1100-1200 series HPLC. Compounds were separated over a Phenomenex Kinetex XB-C18 column (100 × 3 mm, 100 Å, 2.6 μm particle size) held at 50 °C. The mobile phase was composed of 10 mM ammonium formate, pH 4.5 (solvent A) and acetonitrile (solvent B) the following LC method was used:

| Time (min) | %A | %B | Flow rate (µL/min) |
| --- | --- | --- | --- |
| 0 | 98 | 2 | 400 |
| 2 | 98 | 2 | 400 |
| 5.5 | 2 | 98 | 400 |
| 8.5 | 2 | 98 | 400 |
| 9.5 | 98 | 2 | 400 |
| 12.75 | 98 | 2 | 400 |

The mass spectrometer was operated in positive ion mode using the following parameters: Curtain gas: 20 L/min, Collision gas: medium, Ionspray voltage: 4500 V, Temperature: 250 °C, Ion source gas 1: 20 L/min, Ion source gas 2: 10 L/min, Declustering potential: 0, Entrance potential: 10, Collision cell exit potential: 10. Coumaroyl-CoA was detected in multiple reaction monitoring (MRM) mode by monitoring the following transition:914.16 m/z → 407.16 m/z using a collision energy of 47.8 V. Data was imported into Skyline targeted mass spectrometry environment (v. 3.7.0 11317) and peaks were integrated using Skyline’s embedded integration function. Coumaroyl-CoA was quantified relative to a standard curve that was generated by calculating the linear regression of 3 individual injections of each concentration within the standard curve (R^2^ = 0.9982).

## Acknowledgements

We would like to thank Professor Mattheos Koffas for providing us with the plasmid pEMT6-4CL-CUS. We thank Jesus Barajas for his careful reading of this manuscript and helpful suggestions during preparation. We also thank Morgan Price for assistance in analyzing RB-TnSeq data.

This work was part of the DOE Joint BioEnergy Institute (https://www.jbei.org) supported by the U. S. Department of Energy, Office of Science, Office of Biological and Environmental Research, supported by the U.S. Department of Energy, Energy Efficiency and Renewable Energy, Bioenergy Technologies Office, through contract DE-AC02-05CH11231 between Lawrence Berkeley National Laboratory and the U.S. Department of Energy. The views and opinions of the authors expressed herein do not necessarily state or reflect those of the United States Government or any agency thereof. Neither the United States Government nor any agency thereof, nor any of their employees, makes any warranty, expressed or implied, or assumes any legal liability or responsibility for the accuracy, completeness, or usefulness of any information, apparatus, product, or process disclosed, or represents that its use would not infringe privately owned rights. The United States Government retains and the publisher, by accepting the article for publication, acknowledges that the United States Government retains a nonexclusive, paid-up, irrevocable, worldwide license to publish or reproduce the published form of this manuscript, or allow others to do so, for United States Government purposes. The Department of Energy will provide public access to these results of federally sponsored research in accordance with the DOE Public Access Plan (http://energy.gov/downloads/doe-public-access-plan)

## Contributions

Conceptualization, M.R.I., M.G.T., J.M.B; Methodology, M.R.I., M.G.T., J.M.B.,Y.L.,C.J.P.; Investigation, M.R.I., M.G.T., J.M.B., Y.L., M.S., A.N.P., J.W.G.; Writing – Original Draft, M.R.I.; Writing – Review and Editing, All authors.; Resources and supervision, C.J.P, A.M.D., J.D.K.

---

**Figure S1:**
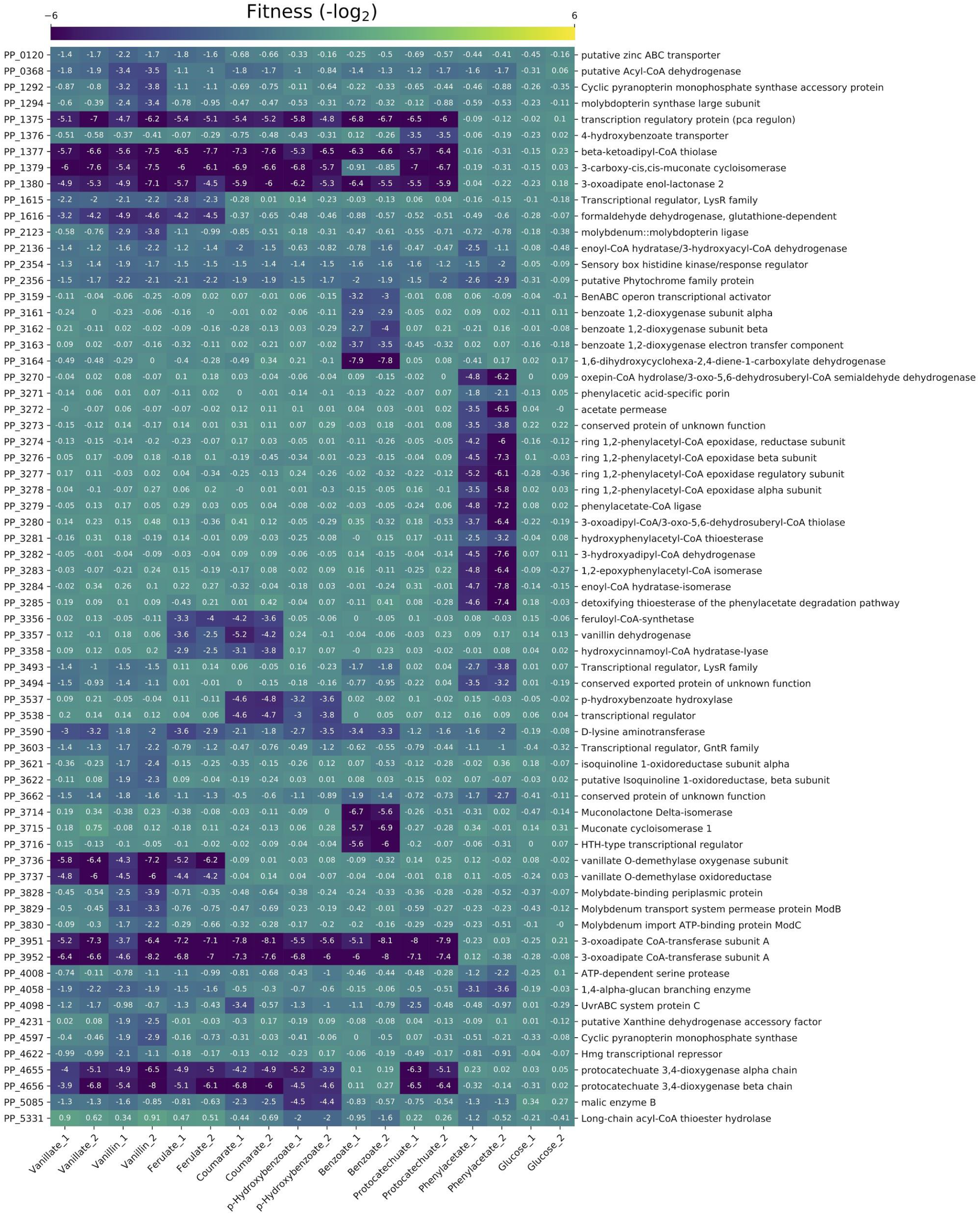
Heatmap of RB-TnSeq mutant fitness data. Genes shown have fitness defects (−log_2_ < −2) and with statistical significance (|t_score_| > 4) in at least one tested condition which show no significant fitness defects when grown on glucose.

**Figure S2:**
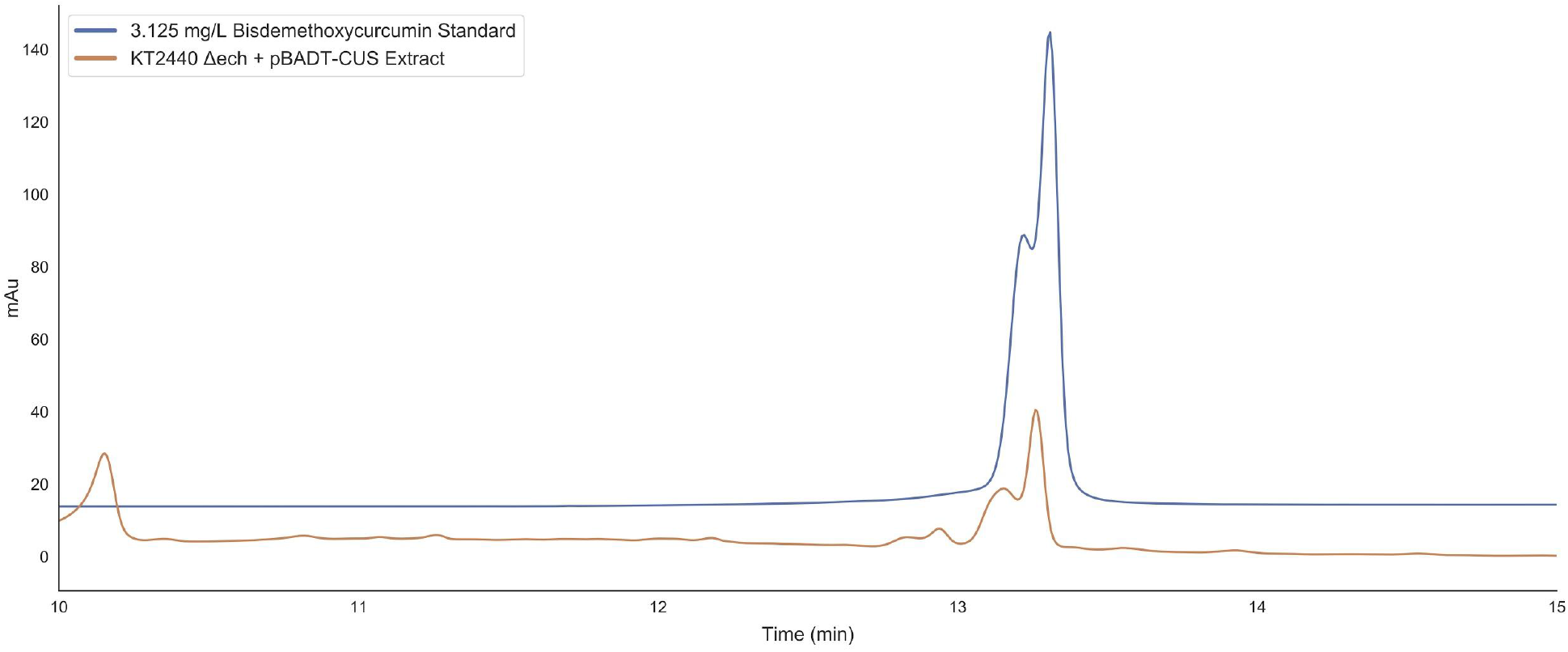
HPLC trace of curcumin extracted from *P. putida* KT2440 Δ*ech* + pBADT-CUS overlaid with a 3.125 mg/L standard

**Figure S3:**
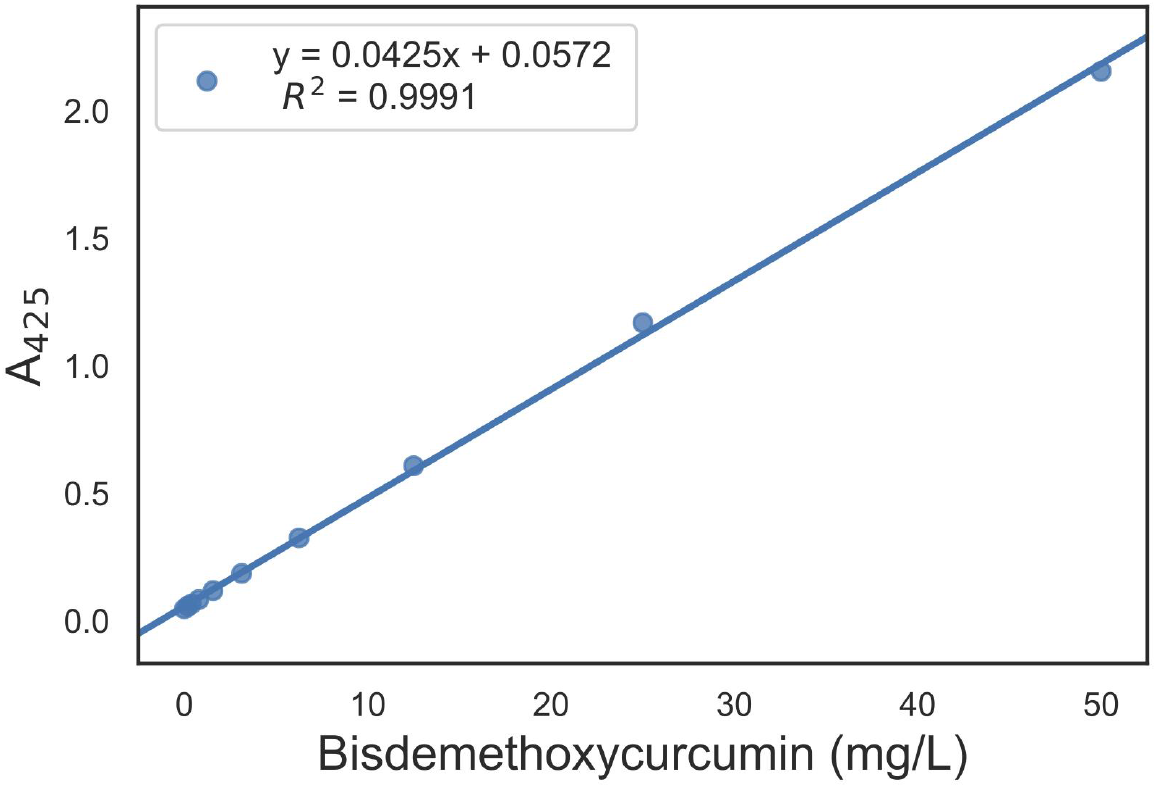
Standard curve of bisdemethoxycurcumin. Absorbance measurements at 425 nm from 50 to 0 mg/L of bisdemethoxycurcumin dissolved in oleyl alcohol.

**Figure S4:**
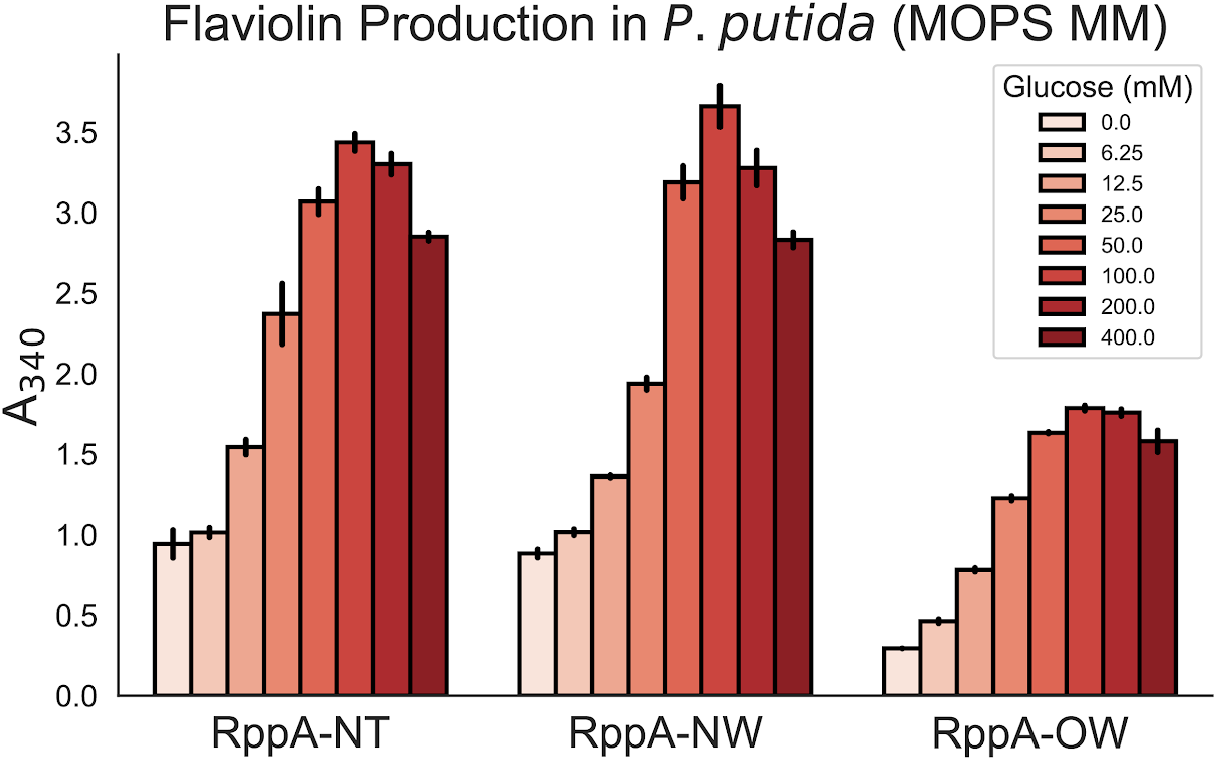
Production of flaviolin in *P. putida* KT2440 measured with absorbance at 340 nm. Glucose concentrations were varied from 0 mM to 400 mM in MOPS minimal medium.

**Figure S5:**
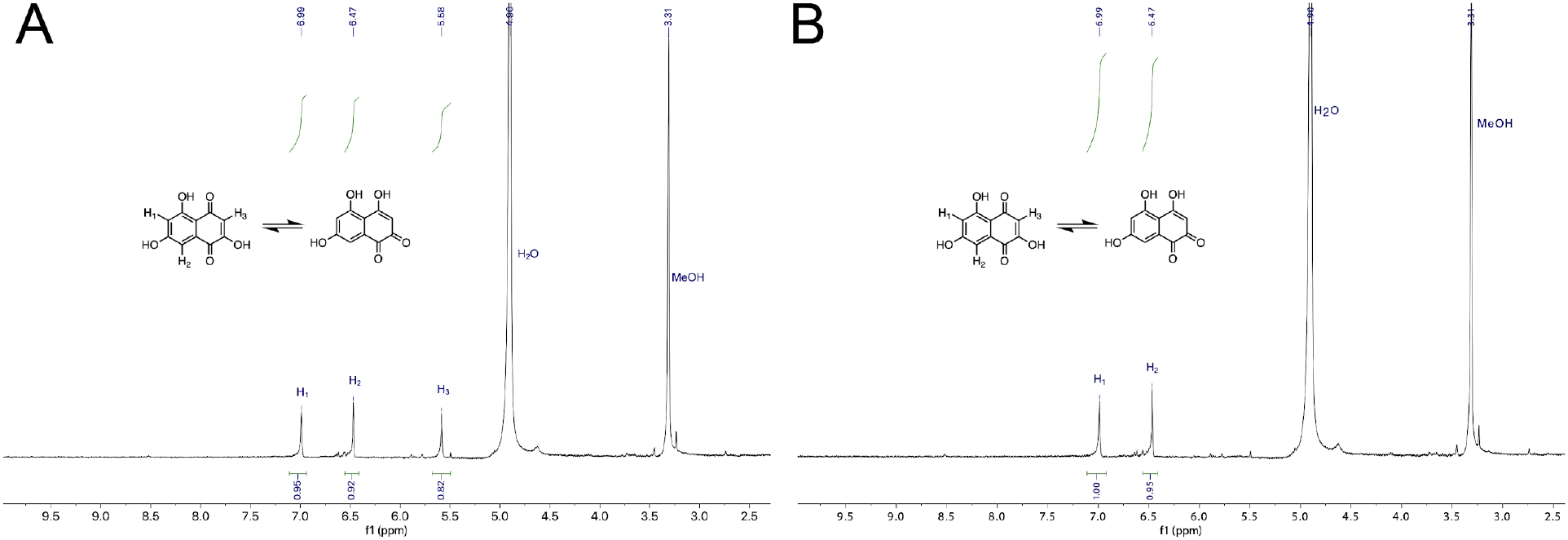
^1^H-NMR spectrum of flaviolin isolated from *P. putida* KT2440 expressing *rppA*-NT from the plasmid pBADT-*rppA*-NT. A) ^1^H-NMR spectrum from flaviolin freshly resuspended in CD_3_OD. B) ^1^H-NMR spectrum from flaviolin resuspended in CD_3_OD and incubated at room temperature overnight. The signal at 5.58 (s, 1 H) is absent in the overnight treatment due to proton exchange with CD_3_OD (Snyder et al., 2009).

**Figure S6:**
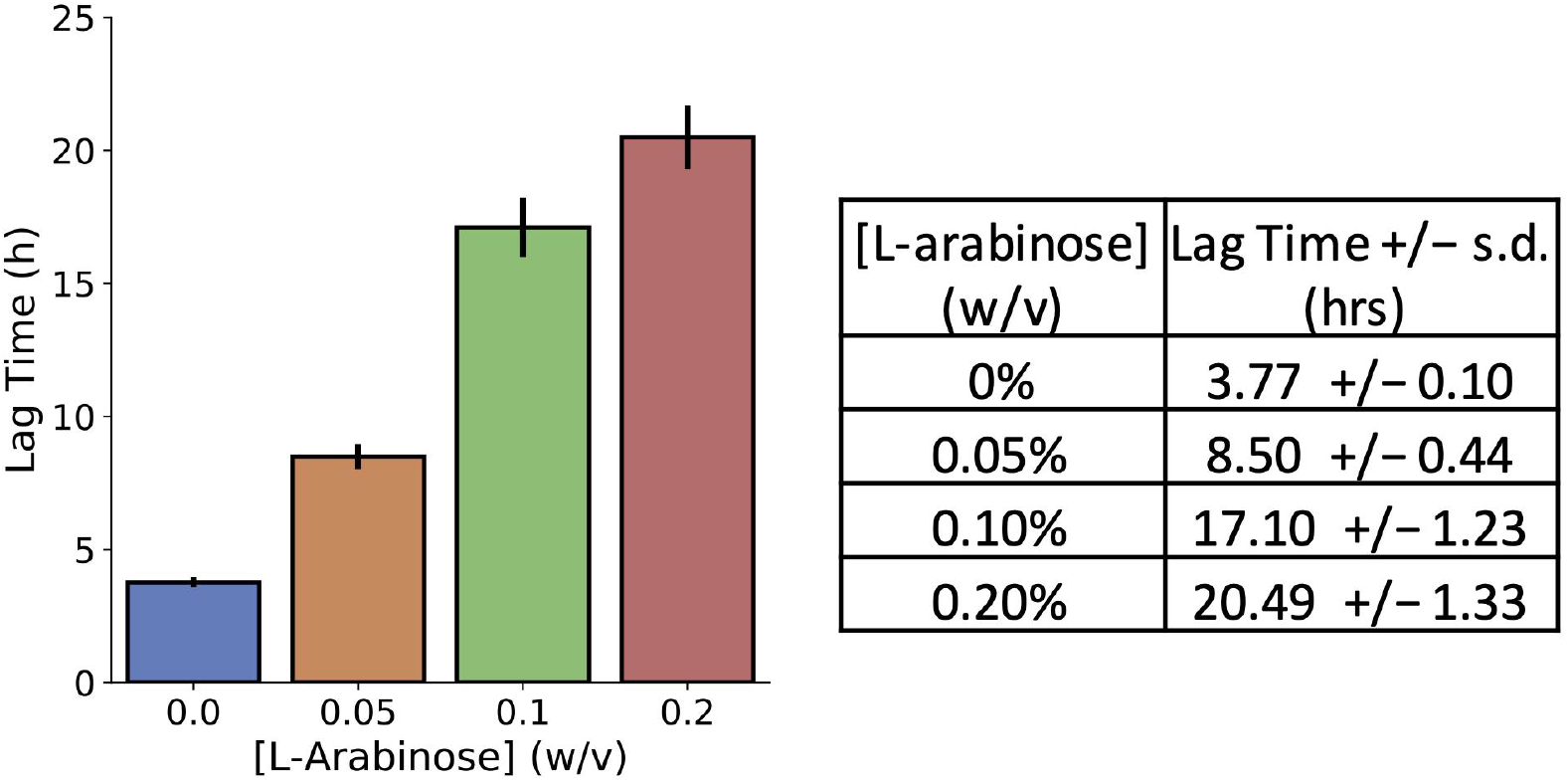
Calculated lagtimes for cultures expressing *fcs* in the presence of 5 mM coumarate. The *fcs* gene was expressed with 0, 0.05, 0.1, or 0.2% w/v L-arabinose. The lagtimes and their standard deviations are included in the table (right). Lag time is defined here as the amount of time required for the culture to reach OD_600_ = 0.16. Standard deviation was calculated from 3 replicates

